# Deciphering the maturation of tertiary lymphoid structures in cancer and inflammatory diseases of the digestive tract using imaging mass cytometry: from high-level data to a simple architectural and functional grading

**DOI:** 10.1101/2022.11.15.516576

**Authors:** Marion Le Rochais, Patrice Hémon, Danivanh Ben-guigui, Soizic Garaud, Christelle Le Dantec, Jacques-Olivier Pers, Arnaud Uguen

## Abstract

**Objective:** Persistent inflammation can promote the development of tertiary lymphoid structures (TLS) within tissues resembling the secondary lymphoid organs (SLO) as lymph nodes (LN). The composition of the TLS across different organs and diseases could be of pathophysiological and medical interest. In this work, we compared TLS to SLO and between cancer and inflammatory diseases of the digestive tract.

**Design:** Colorectal and gastric tissues with different inflammatory diseases and cancers from the department of pathology of CHU Brest were analyzed based on 39 markers using imaging mass cytometry (IMC). Unsupervised and supervised clustering analyses of IMC images were used to compare SLO and TLS.

**Results:** Unsupervised analyses tended to group TLS per patient but not per disease. Supervised analyses of IMC images revealed that LN had a more organized structure than TLS and non-encapsulated SLO Peyer’s patches. TLS followed a maturation spectrum with close correlations between germinal cell (GC) markers’ evolution. The correlations between organizational and functional markers made relevant the previously proposed TLS division into three stages: lymphoid-aggregates (LA) (CD20+CD21-CD23-) had neither organization nor GC functionality, non-GC TLS (CD20+CD21+CD23-) were organized but lacked GC’s functionality and GC-like TLS (CD20+CD21+CD23+) had GC’s organization and functionality. This architectural and functional maturation grading of TLS pointed to differences across diseases.

**Conclusion:** TLS architectural and functional maturation grading is accessible with few markers allowing future diagnostic, prognostic, and predictive studies on the value of TLS grading, quantification and location within pathological tissues in cancers and inflammatory diseases.

**KEY MESSAGES:**

- **What is already known on this topic:** Tertiary lymphoid structures (TLS) arise in organs under various pathological conditions and can be of prognostic significance.
- **What this study adds:** This study deciphers the composition of TLS in digestive cancers and inflammatory diseases using massively multiplexed (39 markers) imaging mass cytometry (IMC). Beyond the term TLS, this study points to the heterogeneity of these structures in terms of composition and maturation but also the relevance of a simple architectural and functional three-stage grading of TLS.
- **How this study might affect research, practice, or policy:** This preliminary study paves the way for future studies evaluating the diagnostic, prognostic and theranostic values of TLS maturation grading, quantification and location within tissues as novel biomarkers in inflammatory diseases and cancers.

## INTRODUCTION

### Tertiary lymphoid structures

Inflammation is the immune system’s response to various diseases, such as cancer, autoimmune diseases, and infections occurring within the secondary lymphoid organs (SLO), particularly in the draining lymph nodes (LN). In case of persistent inflammation, the migration and positioning of immune cells follow the organogenesis of SLO within the organs, to give rise to tertiary lymphoid structures (TLS) detectable as nodular lymphoid-aggregates (LA) at the microscopic level.(1–3) TLS can develop in infectious, autoimmune, and inflammatory diseases, as well as in solid cancers with a potential significance in terms of prognosis and response to treatments.(4,5) Studies on several diseases causing chronic inflammation proposed staging classifications of TLS depending notably on the density of CD20^+^ B cells and the formation of a follicular structure in TLS.(6–8) But the authors have used different structural, cell differentiation, and functional markers to describe various degrees of TLS, often with limited and heterogeneous panels of markers. This methodological heterogeneity in TLS staging is a limitation for the comparisons of data between studies. A clearer and more reproducible definition of what is a TLS and how mature it is regarding its cell composition, organization, and functionality is needed to better investigate the diagnostic, prognostic, and predictive values of TLS analysis.

### TLS as a potential biomarker in gastric and colorectal diseases

TLS are particularly frequent in various digestive tract diseases. As bacterial and viral infections have been reported in the initiation of TLS formation at the site of infection, TLS are notably observed in the stomach in case of *Helicobacter pylori* inducing chronic gastritis.(9) TLS are also observed in chronic inflammation caused by autoimmune diseases (such as Biermer’s disease – autoimmune gastritis) or inflammatory diseases (Crohn’s disease, Chronic diverticulitis, and Ulcerative colitis).(10,11) Their role in inflammation and autoimmunity is difficult to establish: in murine models, TLS could have a dual function, with opposite pathogenic effects.(12) Indeed, TLS could play a role in pathogen clearance beneficial for the patient, as well as being sites of autoimmune response amplifications and tissue damage aggravations leading to a deleterious disease evolution.(13)

In the cancer field, TLS could also play a dichotomous role in tumor invasion control and metastasis. TLS are privileged sites for antigen presentation to B cells at the tumor site, leading to the increase of the local production of antitumor immunoglobulins G, helping in tumor control and improve patient’s prognosis.(13) Contrastingly, TLS can recruit immunosuppressive cells and become immunological micro-niches promoting the generation of progenitor cancer cells leading to deleterious outcomes.(14) Thus, the prognostic significance of TLS remains unclear. Interestingly, in some colorectal and gastric cancers with a deficit in the repair of DNA replication errors (causing a “microsatellite instability” MSI status), the resulting increased number of neoantigens related to numerous mutations gives rise to highly immunogenic tumors, notably rich in TLS. MSI tumors are of different prognosis in comparison with tumors lacking this deficiency (called “stable microsatellites” (MSS)). Contradictory data are reported about the better prognosis of MSI tumors at an early stage but their poor prognosis in patients with advanced/metastatic MSI tumors. Due to their high immunogenicity, the growth of MSI tumors is conditioned by a high level of immunomodulation, targeted by immune checkpoints inhibitors treatments to improve patients’ survival. Furthermore, regardless of patient characteristics, MSI status and the TLS presence generally correlate with improved survival in CRC patients.(7,15,16)

### Studying TLS in gastric and colonic tissues using imaging mass cytometry

The frequency of TLS arising in various gastric and colonic diseases makes stomach and colon predilection sites for a better characterization of TLS. However, the small size and unpredictable location of TLS within the tissues, detectable only during microscopic examination of tissues, are limitations for their detailed analysis. Thus, it requires highly multiplexed methods able to extract in-depth data from minimal tissue areas selected at the histopathological microscopic level. The recent imaging mass cytometry (IMC) technique allows to co-analyze nearly 40 markers concomitantly in the same tissue section with increasing applications especially in cancer research.(17,18) IMC is particularly appropriate to analyze in-depth the phenotypes and spatial organization of cells in small tissular structures such as TLS.

The objectives of the present work were to decipher and compare the compositions and organizations of TLS observed in different inflammatory and cancerous, gastric and colorectal diseases using IMC.

## MATERIAL AND METHODS

### Cases and samples selection

Tissues samples from patients analyzed for care purpose in the department of pathology of CHU Brest were used, following our national and institutional guidelines, in compliance with the Helsinki Declaration and after approval by our institutional review board, all samples were included in the registered tissue collection part of the Brest Biological Resources Center BB-0033-00037, NF S 96-900 certificated Brest, CHRU Brest AC-2019-3642 – DC – 2008 – 214). No clinical, epidemiological or evolution data about the patients was collected during this study. The patient groups with gastric samples were constituted as follows : (A) control patients with gastric weight-reduction surgery with lymphoid islet with healthy mucosa; (B) patients with autoimmune gastritis (Biermer disease); (C) patients with *Helicobacter pylori*–related gastritis; (D) patients with MSI gastric adenocarcinoma differentiating those without nodal metastasis (D1) from those with nodal metastasis (D2); (E) patients with MSS gastric adenocarcinoma without nodal metastasis (E1) or with nodal metastasis (E2). In the same manner for colorectal tissues: (1): control patients with pT0N0 status colectomy specimens complementary to endoscopic resection of pT1 colorectal cancers, Peyer’s patches were selected distant to the resection site next to the resection margins; (2) patients with Crohn’s disease; (3) patients with ulcerative colitis; (4) patients with chronic diverticulitis; (5) patients with MSI colonic adenocarcinoma differentiating those without nodal metastasis (5A) from those with nodal metastasis (5B); (6) patients with MSS colonic adenocarcinoma without nodal metastasis (6A) or with nodal metastasis (6B). For patients with gastric or colonic adenocarcinomas, one regional LN without nodal metastasis was also selected, completed by a LN with nodal metastasis for metastatic patients (**Table 1**). The costs and workflows constraints of IMC technology haves prone us to design a highly qualitative study selecting only small groups of patients with different diseases (3 to 6 patients per group). The formalin-fixed paraffin-embedded (FFPE) tissue samples and corresponding tissue slides were collected for area containing TLS selection on the histopathological slide and IMC analyses on new tissue sections.

**Table 1:** Cases and regions of interest selected for imaging mass cytometry. ROI: region of interest selected for imaging mass cytometry analysis; T: primary tumor site, LN: non-metastatic lymph node, LNm: metastatic lymph node

| Organs | Group | Condition | Cases number | ROI number |
| --- | --- | --- | --- | --- |
| Stomach | **A** | No gastric disease | 6 | 14 |
|  | **B** | Biermer's autoimmune gastritis | 4 | 10 |
|  | **C** | *Helicobacter pylori* –related gastritis | 5 | 8 |
|  | **D** | Microsatellite instable gastric adenocarcinoma | **6** | **45** |
|  | D1 | Non metastatic (nm) | 3 | 18 (11T, 7LN) |
|  | D2 | Metastatic (m) | 3 | 27 (14T, 5LN, 8LNm) |
|  | **E** | Microsatellite stable gastric adenocarcinoma | **6** | **53** |
|  | E1 | Non metastatic (nm) | 3 | 18 (12T, 6 LN) |
|  | E2 | Metastatic (m) | 3 | 35 (21T, 7LN, 7LNm) |
| **TOTAL** |  |  | **27** | **130** |
| Colon | **1** | Peyer's patches | 5 | 11 |
|  | **2** | Crohn's disease | 5 | 19 |
|  | **3** | Ulcerative colitis | 5 | 14 |
|  | **4** | Chronic diverticulitis | 5 | 10 |
|  | **5** | Microsatellite-instable colonic adenocarcinoma | **6** | **55** |
|  | 5A | Non metastatic (nm) | 3 | 25 (18T, 7LN) |
|  | 5B | Metastatic (m) | 3 | 30 (12T, 8LN, 10LNm) |
|  | **6** | Microsatellite-stable colonic adenocarcinoma | **6** | **50** |
|  | 6A | Non metastatic (nm) | 3 | 15 (4T, 11LN) |
|  | 6B | Metastatic (m) | 3 | 35 (10T, 9LN, 16LNm) |
| **TOTAL** |  |  | **32** | **159** |

### Selection of a 39 markers-panel for TLS description

We developed and validated 39 metal-tagged antibodies IMC panel, composed of structural, functional, and immune markers for the in-depth description of TLS. 7 immune markers were selected for T cell populations (CD3/CD4/CD8/Tbet/GATA3/Stat3/FoxP3), 4 immune markers for B cell and plasma cell populations (CD20/CD138/CD38/CD23/CD27), and 9 other markers for DC, macrophages, granulocytes, and NK cells. 5 markers (CD31/CD34/Podoplanin/Pankeratin/CD103) were selected as non-immune markers to enhance visualization of tissue architecture: vessels, epithelium, and fibroblasts. We also selected 13 functional markers (Bcl6/AID/HLADR/CXCR5/CD86/Ki67/Cleaved-Caspase-3/PD-1/PD-L1/CD27/IgM/IgD/IgG) to evaluate the TLS functions in terms of activation, proliferation, inhibition, checkpoint, cytokines and immunoglobulins. A nuclear intercalator dye (Iridium) was included to allow the identification and segmentation of individual cells in our analysis pipeline (**Table 2**).

**Table 2:** Panel of markers and corresponding metals-labeled antibodies for tertiary lymphoid characterization

| Marker | Metal | Supplier | Dilution | Antibody clone | Metal coupling: in-house (I) or manufacturer (M) |
| --- | --- | --- | --- | --- | --- |
| **CD83** | 139La | Abcam | 1/100 | EPR23809-19 | I |
| **CD38** | 141Pr | Fluidigm | 1/100 | EPR4106 | M |
| **CD21** | 142Nd | Abcam | 1/400 | SP186 | I |
| **CD23** | 143Nd | Abcam | 1/500 | EPR3617 | I |
| **DCLAMP** | 144Nd | Novus | 1/50 | 1010E1.01 | I |
| **T-bet** | 145Nd | Fluidigm | 1/50 | D6N8B | M |
| **CD34** | 146Nd | Abcam | 1/200 | SP179 | I |
| **CD163** | 147Sm | Fluidigm | 1/200 | EDHu-1 | M |
| **Pankeratin** | 148Nd | Fluidigm | 1/200 | C11 | M |
| **GATA-3** | 149Sm | Abcam | 1/300 | EPR16651 | I |
| **CD274 (PD-L1)** | 150Nd | Abcam | 1/75 | SP142 | I |
| **CD31** | 151Eu | Fluidigm | 1/100 | EPR3094 | M |
| **Ki-67** | 152Sm | Abcam | 1/300 | MKI67/2642 | I |
| **IgD** | 153Eu | Southern Biotech | 1/40 | Polyclonal | I |
| **AID** | 154Sm | Abcam | 1/500 | EPR23436-45 | I |
| **FoxP3** | 155Gd | Abcam | 1/50 | 236A/E7 | I |
| **CD86** | 156Gd | RD | 1/100 | Polyclonal | I |
| **CD68** | 159Tb | Fluidigm | 1/400 | KP1 | M |
| **Bcl-6** | 160Yb | Abcam | 1/75 | CAL49 | I |
| **CD20** | 161Dy | Fluidigm | 1/400 | H1 | M |
| **CD8a** | 162Dy | Fluidigm | 1/200 | D8A8Y | M |
| **CD138** | 163Dy | Abcam | 1/400 | SP152 | I |
| **MPO** | 164Dy | Abcam | 1/1000 | EPR20257 | I |
| **CD279 (PD-1)** | 165Ho | Abcam | 1/100 | NAT105 | I |
| **CD56** | 166Er | Abcam | 1/50 | NCAM1/1496 | I |
| **Granzyme (GzB)** | 167Er | Fluidigm | 1/200 | EPR20129-217 | M |
| **CXCR5** | 169Er | Abcam | 1/100 | EPR23463-30 | I |
| **CD3** | 170Er | Fluidigm | 1/150 | Polyclonal | M |
| **CD27** | 171Yb | Fluidigm | 1/100 | EPR8569 | M |
| **Caspase-3 cleaved** | 172Yb | Fluidigm | 1/100 | 5A1E | M |
| **Podoplanin** | 173Yb | Abcam | 1/400 | PDPN/1433 | I |
| **HLA-DR** | 174Yb | Fluidigm | 1/600 | YE2/36HLK | M |
| **CD4** | 175Lu | Abcam | 1/300 | EPR6855 | I |
| **IgM** | 176Gd | Dako | 1/100 | Polyclonal | I |
| **Cell intercalator** | 191/193Ir | Fluidigm | 1/500 | - | M |
| **CD11c** | 194Pt | Abcam | 1/300 | ITGAX/1242 | I |
| **CD103** | 195Pt | Abcam | 1/75 | SP301 | I |
| **Stat3** | 196Pt | Abcam | 1/800 | EPR787Y | I |
| **IgG** | 209Bi | Dako | 1/150 | Polyclonal | I |

### Antibodies validation using immunohistochemistry

For the antibodies not already conjugated to metals by the manufacturer, the antibodies’ performances, and optimal antigen retrieval conditions, were assessed by chromogenic immunohistochemistry (IHC). Tissue sections were deparaffinized and rehydrated with xylene and decreasing concentrations of ethanol. Sections were boiled for 20min in Tris-EDTA (10 mM/1 mM, pH 9) buffer for antigen retrieval, followed by endogenous peroxidase blockade using a 0.3% H2O2 solution for 5min and non-specific antibody binding blockade with 3% TBS BSA solution for 30min. Tissues were incubated overnight at 4°C with the primary antibody. Following washes in PBS, tissues were incubated with a secondary antibody coupled to horseradish peroxidase (Polink-1 HRP for Rabbit & Mouse – GBI Labs Kit / AffiniPure Goat Anti-Rat IgG −112-005-143, Jackson / AffiniPure Donkey Anti-Goat IgG – 705-005-003, Jackson) for 1h at room temperature. Antibody binding was detected with DAB+ chromogen (DAKO, Agilent technologies, Santa Clara, Ca, USA) and the sections were counterstained with hematoxylin (Thermo Fisher Scientific, Waltham, Massachusetts, USA), before dehydration and mounting of the slides. The specificities and intensities of the immunostaining are assessed by a pathologist.

### Antibodies and metal conjugation

IHC-validated antibodies were conjugated to purified lanthanide metals (Fluidigm, San Francisco, CA, USA) (**Table 2**) using the MaxPar antibody labeling kit (Fluidigm) according to the manufacturers’ instructions. After conjugation, all coupled antibodies were eluted in antibody stabilizer buffer (Candor Bioscience, Wangen, Germany). The titration of the coupled antibody in IMC was done on the positive tissue to determine the optimal concentration. Three different concentrations, were tested (0.5X, 1X, and 2X where 1X was defined according to the antibody’s concentration in IHC). The best concentration was then chosen to have the highest staining intensity with the less noise background.

### Tissue labeling before imaging mass cytometry acquisition

For downstream analyses, all tissues were cut at 3 μm and placed on Superfrost® Plus slides (Thermo Scientific, Saint-Herblain, France). All slides were stained with alcian blue to record the microscopic morphology of the. Slides were mounted for scanning at 20X magnification on the 3DHistech Panoramic Midi slide scanner (3DHISTECH, Budapest, Hungary), and visualized using CaseViewer software(v.2.4/32bits, 3DHistech) for the selection of ROIs (**Supplementary Figure 1**). With successive xylene baths, coverslips were removed and the mounting medium was washed. The same IHC-protocol was followed except the peroxidase blockade step, before the incubation of slides with the 39 metal-conjugated antibodies panel and the cell intercalator.

### Imaging mass cytometry acquisition

Before the acquisition, the Hyperion mass cytometry system (Fluidigm) was autotuned using a 3-element tuning slide (Fluidigm) according to the tuning protocol provided by Fluidigm. ROIs with sizes ranging from 300 x 300 μm to 1,200 x 1,200 μm were ablated and acquired at 200 Hz. For each tissue, from one to several ROI were defined for the acquisition on Hyperion, depending on the number of TLS on slides.

### Image analysis pipeline based on QuPath

Generated data were visualized as MCD files using the Fluidigm MCD^TM^ viewer. To better separate antibodies signals and noises, each marker was separately visualized and a minimum signal threshold was set in the Fluidigm MCD^TM^ viewer. For each recorded ROI, a stack of 16-bit single-channel OME-TIFF files was exported from MCD binary files using MCD^TM^ Viewer 1.0 (Fluidigm). OME-TIFF files of each marker were then stacked in one TIFF file with ImageJ (v.1.8.0_172). The TIFF files were opened with QuPath (v.0.3.2), setting the image type on fluorescence, for running the segmentation with the best parameters for each ROI, based on the nuclei-staining Iridium channel and peri-nuclear cell expansion.(19) Measurements (mean of the marker signal in each cell) were then exported as a CSV table for analysis.

### Unsupervised and supervised clustering analyses

The unsupervised clustering was made with Cytofkit, a Bioconductor package. The cellular subsets detection was realized by the clustering algorithm PhenoGraph (nearest neighbors k=120). Cytofkit is implemented in R, licensed under the Artistic license 2.0, and freely available from the Bioconductor website, https://bioconductor.org/packages/cytofkit/. The supervised clustering was based on the 15 following markers: CD103, CD138, CD163, CD20, CD27, CD3, CD34, CD4, CD56, CD68, CD8, DCLAMP, MPO, PanKeratin, Podoplanin. No transformation method was used, and a t-sne dimension reduction was applied.

### Pathologic assessment of IMC images

A semi-quantitative supervised analysis was performed with the visual grading of IMC images through 15 markers characteristics of the germinal center (GC) (DCLAMP/CD3/CD8/CD20/CD21/CD23/CD4/ IgD/CXCR5/CD38/Podoplanin/AID/Bcl6/CD68/CD138). The scoring was performed by a pathology-trained scientist blinded to the diagnostic and experimental data. Each marker was analyzed for each TLS and scored from 0 to 4 according to the organization: 0 being the absence of staining and 4 the higher grade of organization, with the aspect very close to a functional GC as expected in LN (**Table 3 and Supplementary Figure 2**). Correlations scores between markers were determined. If the p-value was significant (p<0.05), we considered, as low positive correlations scores higher than 0.5, and as high positive correlations scores higher than 0.7. Statistics and heat maps were generated in R (v4.0.5) with the ggplot2 package.

**Table 3:**
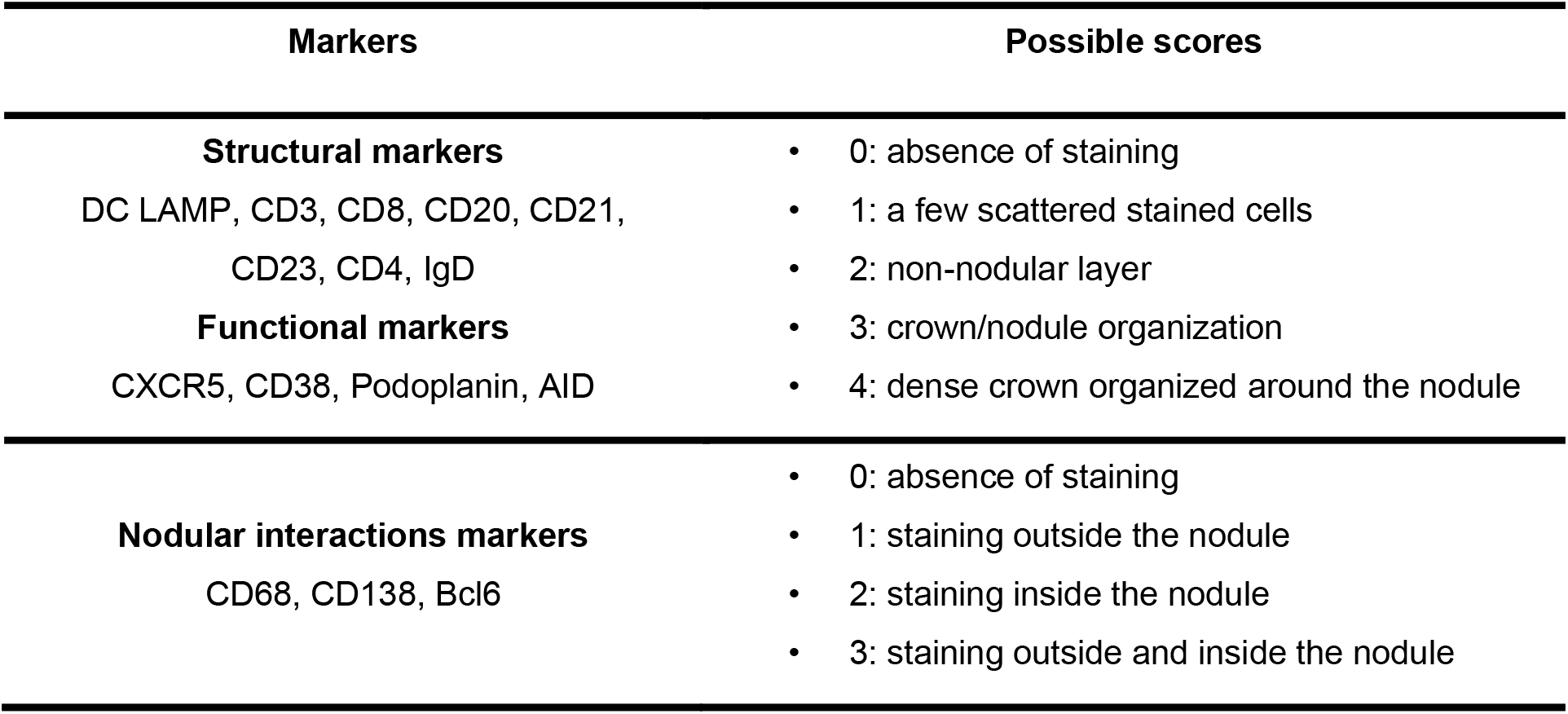
Criteria used for the pathological assessment of germinal center markers on imaging mass cytometry images

In addition, TLS were classed according to the 3 stages classification in terms of LA (CD20+CD21-CD23-TLS), Non-GC TLS (CD20+CD21+CD23-) and GC-like TLS (CD20+CD21+CD23+). Distributions between groups were compared using Kruskal-Wallis non-parametric test performed using Medcalc software.

## RESULTS

### Cases and ROIs selections

The study included a total of 27 cases for gastric samples and 32 cases for colonic samples and 130 ROIs in gastric samples and 159 ROIs in colonic samples (**Table 1**).

### Unsupervised single-cell analysis reveals a patient-dependent TLS clustering

Unsupervised analysis was run to determine if there were similar cellular clusters between the TLS of the same organ, the same disease, or a group of affection (infectious, inflammatory, cancer). The RPhenograph analysis was applied to all ROI (excepting those concerning LN, 90 gastric ROI and 98 colonic ROI) to identify meta-clusters associated with distinct subpopulations and the dendrogram enabled to group TLS expressing same cellular clusters.

This identified 25 distinct meta-clusters for the stomach (**Figure 1A**) and 39 for the colon (**Figure 1B**) enabling the visualization of a heatmap plot with its dendrogram where TLS from the same condition were not grouped. TLS cellular composition appeared to be more patient-dependent, rather than disease-dependent because no cluster was found more specifically in one disease or another. For gastric samples, a majority of TLS from the same sample and patient were grouped within the dendrogram (55/90 (61%) TLS, **Figure 1C**). We found the same for colonic samples with a majority of TLS from the same patient grouped within the dendrogram (60/98 (61%) TLS, **Figure 1D**). Given this analysis, TLS cellular compositions seem to be not dependent on the disease or the organ but could be more dependent on the particular features of each patient’s immune system and proper pathology.

**Figure 1:**
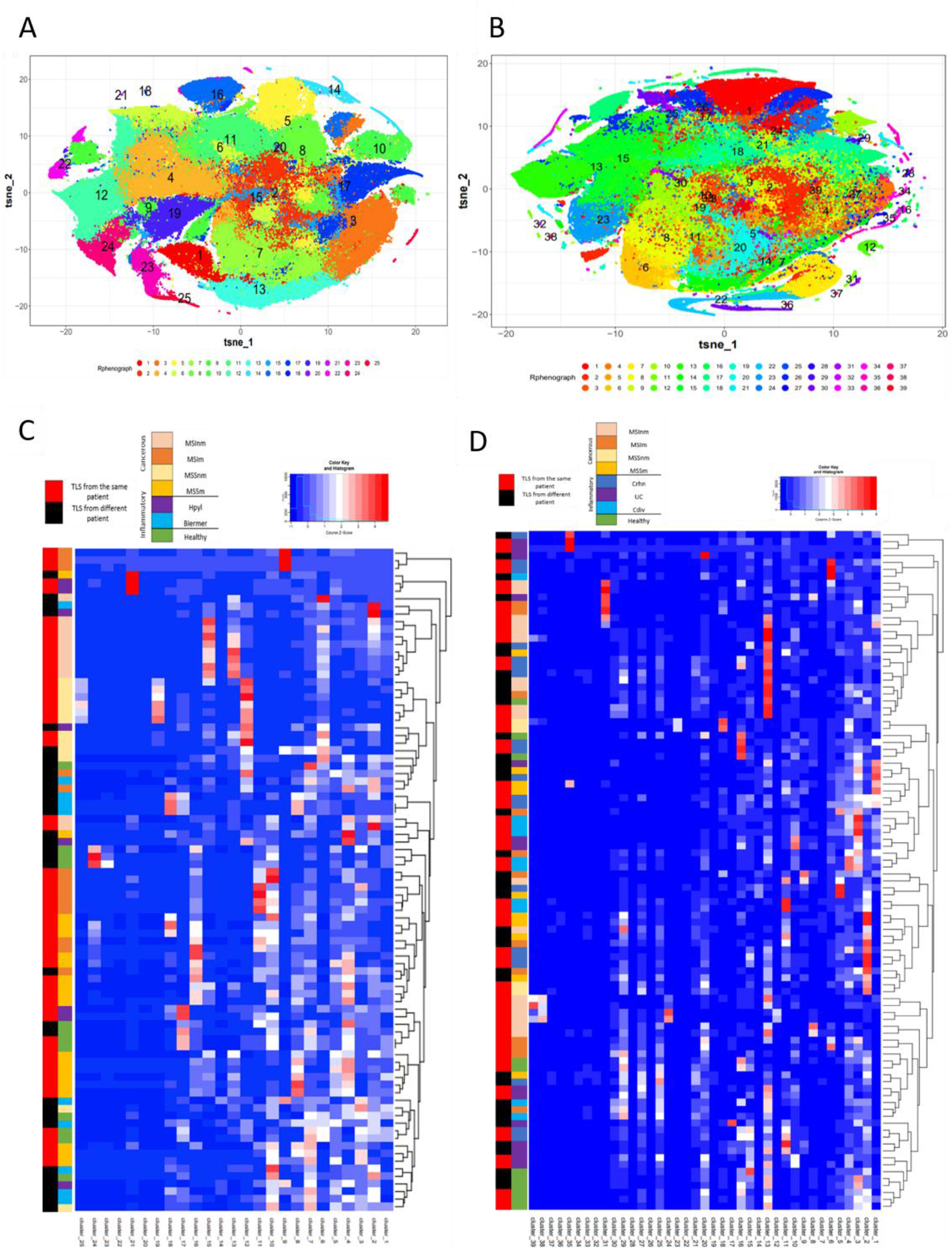
T-sne plots and the corresponding heatmaps of the cellular clusters obtained with Rphenograph composing TLS in gastric and colonic samples. **A**) T-Sne plot showing the cellular composition of each TLS in gastric samples (excluding LN) composed of 25 clusters. **B**) T-Sne plot showing the cellular composition of each TLS (excluding LN) composed of 39 clusters in colonic samples. **C**) Heatmap about gastric data shows median marker expression of clusters detected by RPhenoGraph in TLS coming from the different conditions 1) healthy, 2) inflammatory: Hpyl = patient with infection by *Helicobacter* pylori, Bierm = patient with Biermer’s disease, and 3) cancerous: patients with adenocarcinoma of MSI- or MSS- status, with a non-metastatic (-nm) or metastatic stage (-m). **D**) Heatmap about colonic data shows median marker expression of clusters detected by RPhenoGraph in TLS coming from the different conditions 1) healthy, 2) inflammatory: Crhn = patient with Crohn’s disease, UC = patient with ulcerative colitis, and Cdiv = patient with chronic diverticulitis, and 3) cancerous: patients with adenocarcinoma of MSI- or MSS-status, with a non-metastatic (-nm) or metastatic stage (-m). For C and D heatmaps, row labels represent the cluster IDs and column labels show the TLS name. The multicolor line on the left represents the grouping of TLS according to their conditions where each color is specific for a condition. The red and black line on the left represents in red the TLS that are closely grouped within the dendrogram and found in the same patient whereas in black are TLS from different patients.

### Semi-quantitative analysis based on 15 GC markers highlights a structural organization more important in LN than in TLS

To better understand the development of the GC in TLS and make a comparison with the classical GC found in LN, we made a pathologic assessment of IMC images. We decided to grade 15 chosen markers characterizing GC organization and functions: CD20/CD21/CD23/IgD/CD3/CD4/CD8/DCLAMP/CXCR5 /Podoplanin/CD38/AID/Bcl6/CD138/CD68.

In the stomach, for the 8 structural markers, a difference can be observed between the lymphoid follicles from the LN and TLS from cancerous tissues as well as with TLS from the inflammatory conditions and controls. The first have higher scores (majority of 3 to 4) for the structural markers than the others (**Figure 2A**). The same trend is observed in the colon, with a difference between the LN and TLS from the different conditions (**Figure 2B**), with a higher score for LN independently from their status (metastatic or not) than in TLS, highlighting that LN are structurally better organized than TLS. Of note, SLO Peyer’s patches were closer to TLS than to other SLO, consisting in LN through this analysis.

**Figure 2:**
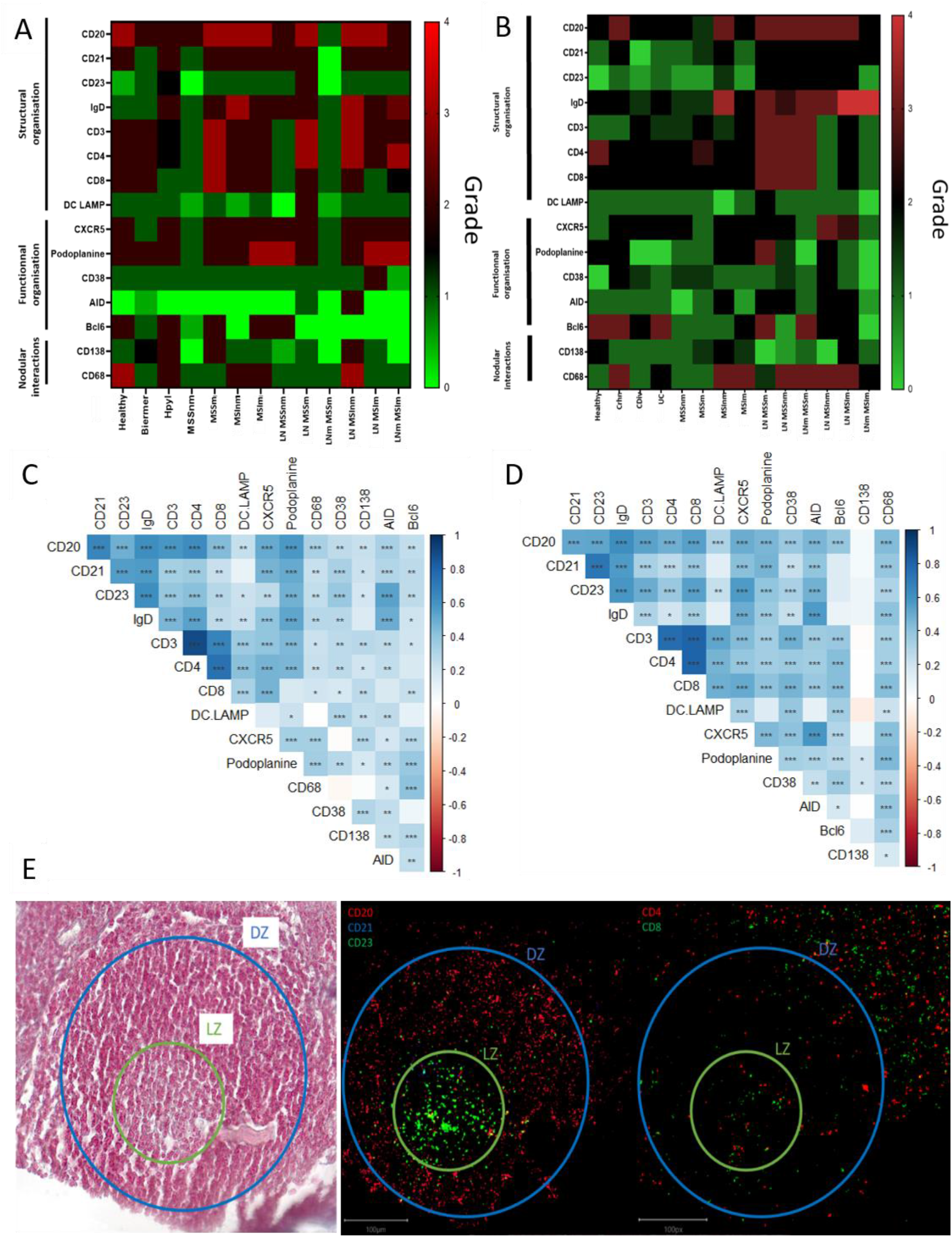
Semi-quantitative supervised analysis based on the scoring of 15 markers characterizing germinal centers and correlations between these markers in the TLS from gastric and colonic samples. **A)** Heatmap representation of the average score observed for each marker for the different conditions observed in the stomach: 1) healthy, 2) inflammatory: Hpyl = patient with infection by *Helicobacter* pylori, Bierm = patient with Biermer’s disease, and 3) cancerous: patients with adenocarcinoma of MSI- or MSS-status, with a non-metastatic (-nm) or metastatic stage (-m). **B)** Heatmap representation of the average score observed for each marker for the different conditions observed in the colon: 1) healthy, 2) inflammatory: Crhn = patient with Crohn’s disease, UC = patient with ulcerative colitis, and Cdiv = patient with chronic diverticulitis, and 3) cancerous: patients with adenocarcinoma of MSI- or MSS-status, with a non-metastatic (-nm) or metastatic stage (-m). The follicles from LN without nodal metastasis (LN) and with nodal metastasis (LNm) are also included in the analysis revealing a more mature structural organization in LN than in TLS. **C)** Heatmap representation of the correlation between the markers in gastric TLS. **D)** Heatmap representation of the correlation between the markers in colonic TLS. Correlated and uncorrelated markers are shown in dark blue to red respectively. Significative p-values are represented with stars: * = p<0.05, ** = p<0.01, *** = p< 0.001. **E)** IMC images of the light (LZ) and dark zone (DZ) in a GC of a TLS from a Crohn’s disease sample with the corresponding HE image on the left.

### Semi-quantitative analysis on non-grouped TLS scoring emphasizes two different areas in TLS

To understand markers associated with the TLS organization, we assessed the correlation between the 15 markers in individual TLS (**Figure 2C and 2D**).

In the stomach (**Figure 2C**), the correlated markers were the T-cells markers such as CD3/CD4 (0.898) and CD4/CD8 (0.746), structural markers of the GC (CD20/CD21/CD23/IgD) and functional ones (podoplanin/AID/CXCR5) (**Figure 2C and Supplementary Figure 3A**). The dendrogram grouped the positive correlations occurring between the marker of T cells (CD3, CD4, CD8) and DCLAMP, markers found in the T-zone of the TLS. We also noted a correlation with the functional marker CXCR5, suggesting the presence of Tfh (T follicular helper) and Tfr (T follicular regulator). In another branch of the dendrogram, markers of B cells and heart of the GC (B-zone): CD20, CD21, CD23, IgD and podoplanin were grouped and presented a low positive correlation (**Figure 2C**).

In the colon, the same trends were observed. The dendrogram grouped positive correlations occurring between organizational markers of the T cell zone (CD3/CD4/CD8/CD38/DCLAMP) and the functional marker Bcl6. We had another part of markers correlated representing more the B-zone with organizational markers (IgD/CD21/CD20/CD23/CD68), or functional markers such as AID and CXCR5. The marker CD138 did not appear to correlate with either of these two groups (T-zone and B-zone) but still appeared to correlate with CD38 and podoplanin (**Figure 2D** and **Supplementary Figure 3B**).

In this manner, for both organs, the same correlated structural markers are observed with two different groups which can be differentiated in B- and T-zone in the TLS composition. The B-zone corresponds to the light zone (LZ) and the dark zone (DZ) of the GC. The LZ consisted of the FDC network with or without the active GC (CD23+) depending on the maturation stage of the TLS. The dark zone corresponding to the non-proliferative B cells (CD20+ CD23-) which are surrounded by T cells (CD4+ and CD8+ ones) (**Figure 2E**). The T-zone contained mature DCLAMP+ dendritic cells (DC), adjacent to a B-zone which included a typical GC comprising B cells embedded within a network of follicular DC.

### Functional description of E-TLS, PFL-TLS and SFL-TLS and quantifications across gastric and colonic diseases

Beyond globally a less-organized structure of TLS than LN and correlations in terms of markers of the light and the dark zones within TLS, the morphological analysis of IMC images in our study revealed heterogeneous organizations ranging from poorly-organized LA to well-organized structures forming follicles or even functional GC in different organs and diseases. Classifying the TLS into three stages lymphoid-aggregates (LA), Non-GC TLS and germinative center GC-like TLS, we observed the following correlations:

Dense LA, corresponding to the E-TLS stage from Silina et al.,(8) lacking CD21+ FDCs and CD23+ GC which were disorganized LA with T cells (CD3^+^CD4^+^ or CD3^+^CD8^+^) and B cells (CD20^+^), mature DC (DCLAMP^+^) and some macrophages (CD68^+^) surrounding lymphatic (podoplanin^+^) and blood vessel (CD31^+^CD34^+^) (**Figures 3A, 3B and 3C**).

**Figure 3:**
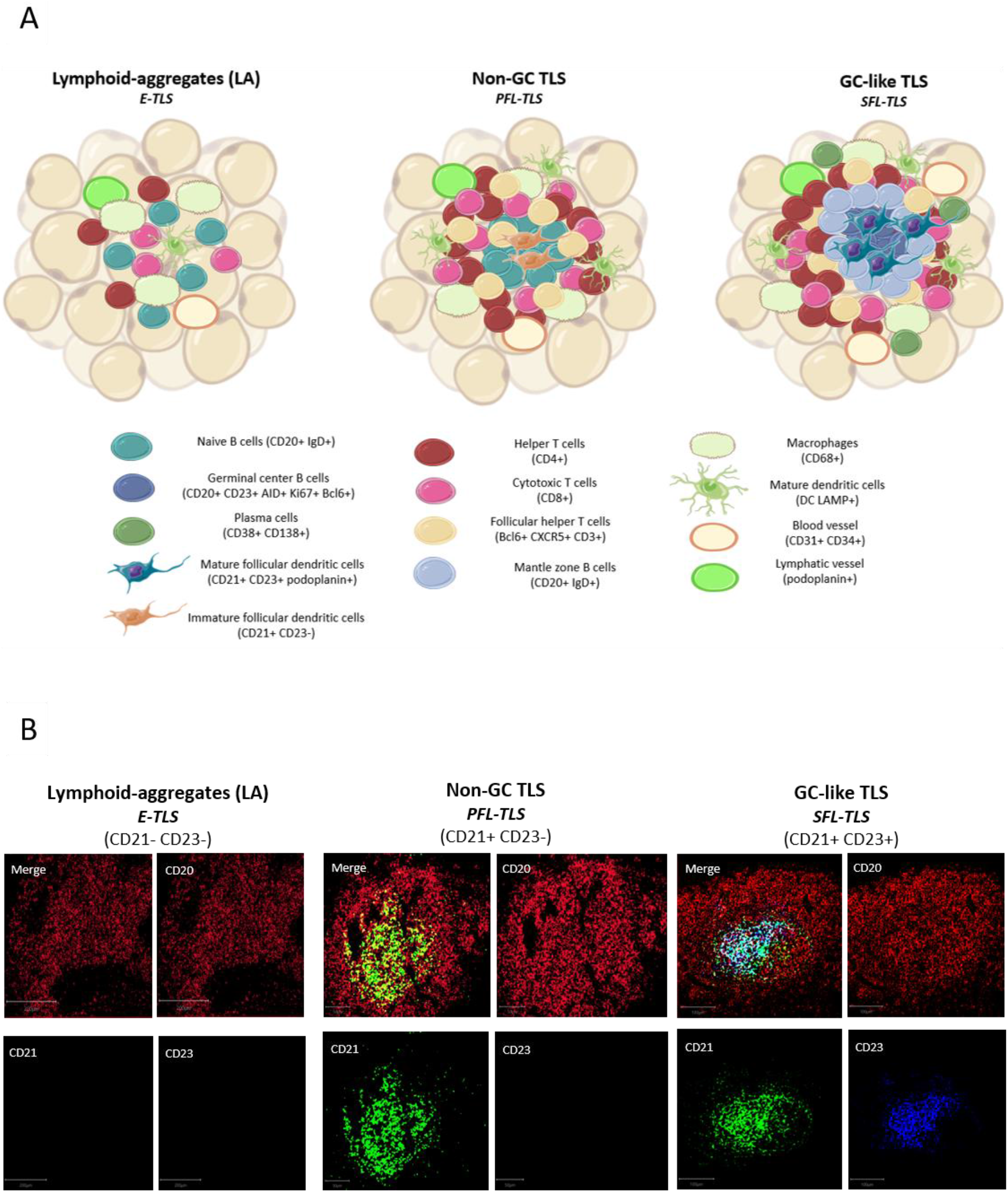
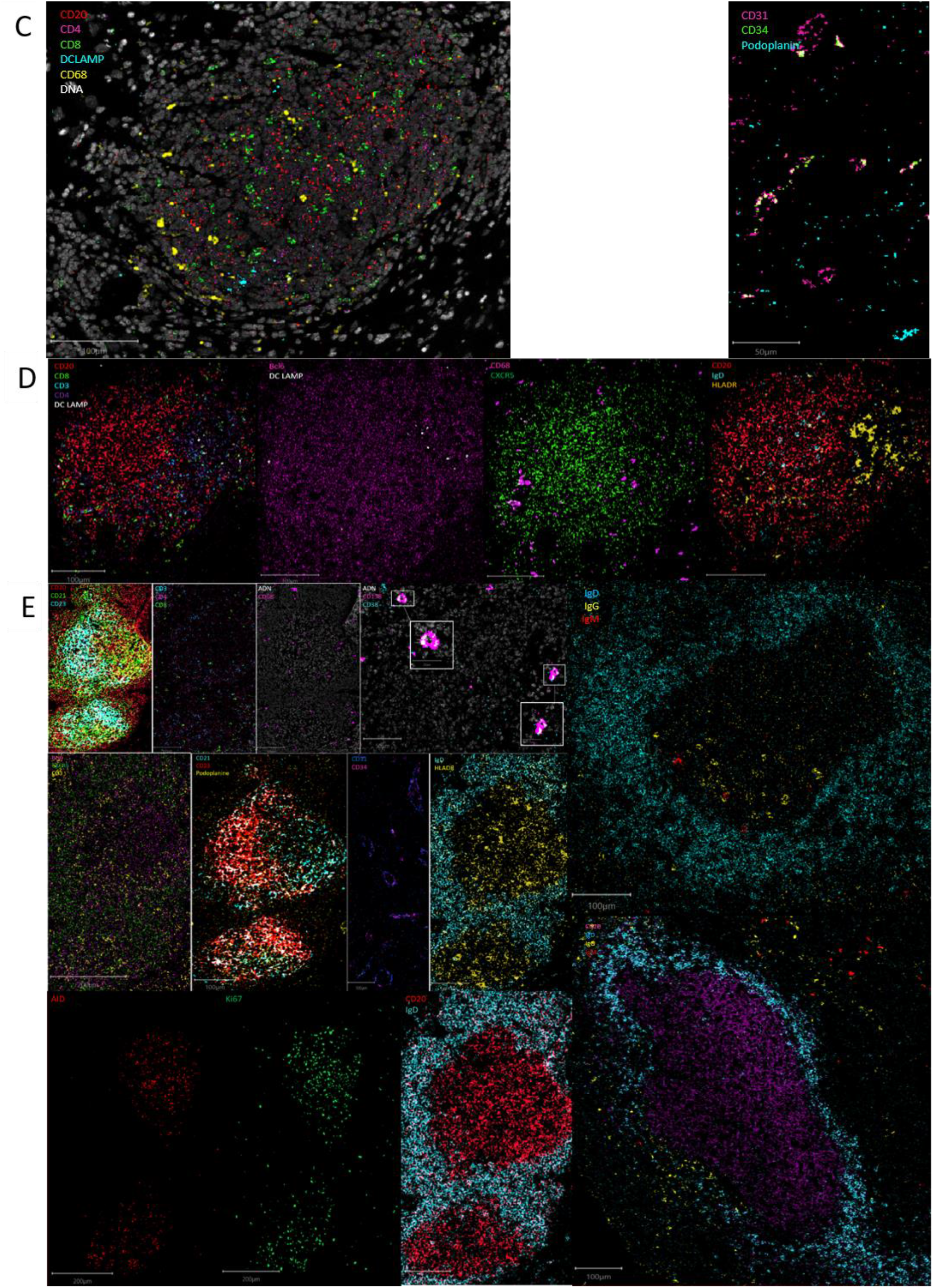
Maturation stages of tertiary lymphoid structures. **A**) The evolutive definition of tertiary lymphoid structures. Lymphoid-aggregates (LA) constitute of T and B cells, with some macrophages and mature dendritic cells, corresponding to early-TLS stage in Silina’s work. Non-germinal center (GC) TLS, corresponding to primary follicle-like TLS, is formed with distinct organized T and B cells zones including follicular dendritic cells in the B-zone. Germinal center (GC)-like TLS, corresponding to secondary follicle-like TLS, is a functional germinal center composed of proliferating mature germinal center B lymphocytes (AID^+^Bcl6^+^Ki67^+^) and follicular dendritic cells (podoplanin^+^) surrounded by mantle zone B cells (IgD^+^) and bordered by follicular helper T cells (Bcl6^+^CXCR5^+^CD3^+^). In addition, TLS are formed of CD4^+^ and CD8^+^ T cells, plasma cells, mature dendritic cells, and macrophages enriched in the T-zone. **B**) Representative images of the IMC markers CD20, CD21 and CD23 characteristics of the three different TLS-stages. Images from the three TLS come from the same Crohn’s patient. **C**) Representative image of IMC markers found in immune aggregate of a patient with Biermer disease. **D**) Representative images of IMC markers found in non-GC TLS of a Crohn’s patient. **E**) Representative images of IMC markers found in GC-like TLS found in one colonic MSI adenocarcinoma and two gastric MSS samples. White boxes show the plasma cells (CD38^+^CD138^+^) appearing as a colocalization of a cyan and a pink signal.

Non-GC TLS defined as LA with immature CD21+ FDCs but no CD23+ GC reaction, corresponding to the PFL-TLS stage.(8) They had a nodular organization with a B cell nodule composed of naïve B cells (IgD+CD20^+^) and follicular helper T cells (Bcl6+CXCR5+CD3+). The B cell zone was surrounded by a T cell zone organized as a crown of helper (CD4^+^) and cytotoxic T cells (CD8^+^) in interaction with some mature DC (DCLAMP^+^) and macrophages (CD68^+^). These non-GC TLS also contained HLADR+ cells allowing antigen presentation (**Figures 3A, 3B and 3D**).

GC-like TLS are defined as LA with mature FDCs (CD21+CD23+) and a functional and organized GC, corresponding to the PFL-TLS stage.(8) This GC had proliferative (Ki67^+^) GC B cells (CD23^+^) expressing AID, involved in somatic hypermutations and class switch recombination, as well as Bcl6+ cells, Bcl6 being a transcription factor involved in GC B cell maturation and follicular T cell differentiation. In addition to FDCs (CD21^+^CD23^+^podoplanin^+^), the proliferative center was surrounded by mantle zone activated B cells (IgD^+^CD20^+^) and contained some macrophages (CD68^+^) as well as CD4^+^ and CD8^+^ T cells (CD3^+^). We also observed a crown of follicular helper T cells (Bcl6^+^CXCR5^+^CD3^+^) and plasma cells (CD38^+^CD138^+^) often surrounding GC indicative of an ongoing humoral and cytotoxic immune response, by the production of IgG and IgA (**Figure 3A, 3B and 3E**).

This classification better suits to the definition of TLS, i.e an organized aggregate of immune cells, where the first stage called LA, corresponding to the E-TLS stage in Silina’s work,(8) is not already considered as a TLS, since nothing can predict the evolution of the disorganized LA in an organized structure meeting the definition of TLS.

Among the different TLS from the same patient, we observed the co-existence of the three stages of TLS evolution (**Figure 3A**), even with the 3-markers-based classification (**Figure 3B**). We also observed significantly different distributions of these three stages between different gastric and colonic pathological sample groups (**Figure 4**) (Kruskal-Wallis test, p-value=0.000115). E.g. gastric MSS adenocarcinomas with regional nodal metastatic spreading had more Gc-like TLS whereas LA were predominant in gastric MSS adenocarcinomas without regional metastasis.

**Figure 4:**
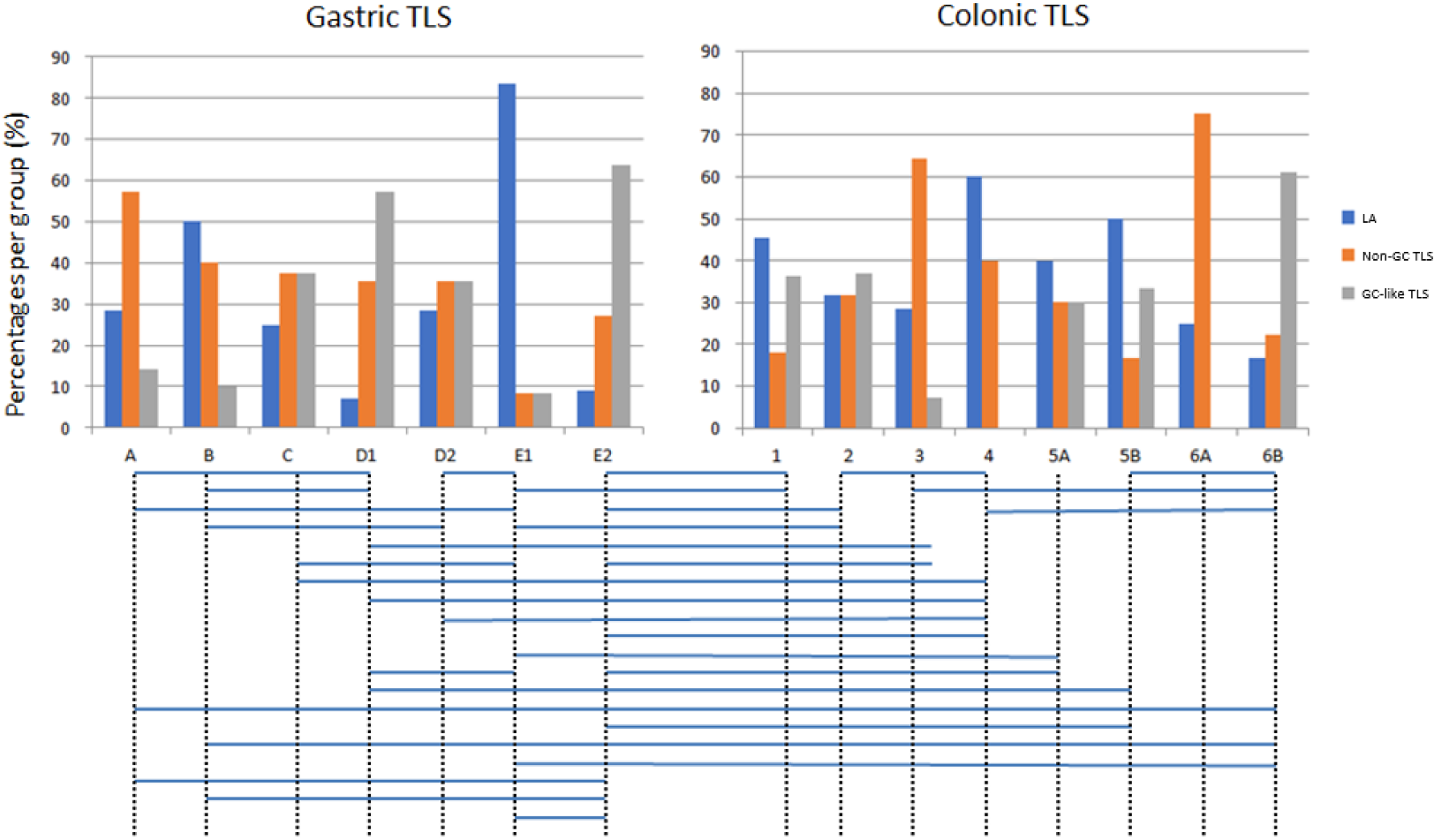
Distributions of the three stages of TLS across the different sample groups. Groups with significantly different distributions (p<0.05) are joined by horizontal bars under the histograms. LA: lymphoid aggregates; Non-GC TLS: non-germinative center TLS; GC-like TLS: germinative center-like TLS; groups A to E2 correspond to gastric samples: A: no gastric disease, B: Biermer’s autoimmune gastritis, C: *Helicobacter pylori*-related gastritis, D1: microsatellite instable gastric cancers without nodal metastasis, D2: microsatellite instable gastric cancers with nodal metastasis, E1: microsatellite stable gastric cancers without nodal metastasis, E2 microsatellite stable gastric cancers with nodal metastasis; groups 1 to 6B correspond to colonic samples: 1: Peyer’s patches, 2: Crohn’s disease, 3: ulcerative colitis, 4: chronic diverticulitis, 5A: microsatellite instable colonic cancers without nodal metastasis, 5B: microsatellite instable colonic cancers with nodal metastasis, 6A: microsatellite stable colonic cancers without nodal metastasis, 6B: microsatellite stable colonic cancers with nodal metastasis.

## DISCUSSION

In recent years, TLS have been studied and described in different tissues and organs, being the subject of a growing number of publications, underlining the evidence that TLS can consist of valuable prognostic and predictive biomarkers to anticipate the aggressiveness of a cancer and responses to anti-cancer treatments including immune checkpoint inhibitors.(5,20–26) To date, TLS are not quantified nor qualified in the pathological examination and, despite their well-demonstrated relation to diseases’ evolutions, they are not taken into account for any treatment decisions. Before TLS characterization could be assessed by pathologists for a treatment-contributive purpose, several points have to be addressed.

Our study was the first to use IMC to concomitantly analyze 39 markers about the different cell types but also functions related to TLS to answer questions related to the composition and organization of TLS in various digestive diseases, gastric and colonic ones, cancerous and non-cancerous ones. Through this study, unsupervised analyses of the 39 markers concluded that TLS compositions were not specific to a given disease but the more pronounced clustering of TLS phenotypes per patient supports that TLS compositions may reflect each patient’s proper immune reaction to a given pathological process.

Comparisons in terms of cell contents and organizations between SLO and TLS confirmed that ectopic TLS remained less organized than SLO. Of note, based on our scoring method, Peyer’s patches (that are considered as SLO and not TLS) seemed to be closer to TLS than LN in terms of organization; this could emphasize the role of the capsule in the superior organization of LN in comparison with non-encapsulated and less organized lymphoid structures as Peyer’s patches and TLS. Nevertheless, correlations between markers were consistent with a progressive organization through a maturation process from non-organized and non-functional TLS to highly organized and functional ones. This is concordant with the maturation of TLS reported in the literature towards several different sets of markers, most of them being included in our IMC TLS-dedicated panel. For future clinical applications it will not be feasible to assess the maturation of TLS through highly-multiplexed but also money- and time-consuming analyses as IMC. So, few markers and appropriate inter-grade boundaries are needed to permit the grading of TLS through routine pathological methods. As proposed by Silina and Posh, TLS in our series could be separated into our three categories only on the basis of three markers CD20, CD21 and CD23 for the characterization of immatures (CD21^+^) and matures FDCs (CD21^+^CD23^+^) and GC B cells (CD20^+^)(**Figure 3B**).(5–8) Towards this three-stages classification, we observed inter-organs and inter-diseases significant differences in the maturation degree of TLS. The small number of TLS and samples in various diseases analyzed through IMC in our study prevents us to draw formal conclusions and additional work with larger numbers of TLS and samples in series of patients with clinical data related to diseases evolution and responses to treatments will be necessary to investigate whether this maturation classification of TLS could consist in novel clinically relevant biomarkers. Nonetheless, this 3-markers approach, correlating with the evolution of other GC IMC markers in our study, appears as a valuable way to class TLS and would be easy to apply through routine pathology methods as multicolor-IHC. This ship towards IHC will also have the advantage to allow analyses of whole tissue slide images (WSI), hardly feasible for cost and technical reasons in IMC. These WSI analyses will provide additional data about TLS, notably in cancer tissues on the locations and counts of TLS.

## CONCLUSION

The complexity and heterogeneity of TLS between organs and diseases are real at the level of cell compositions and interactions as underlined by this study using in-depth highly multiplexed IMC. Nevertheless, correlations between markers used to decipher this complexity permit to draw a more simple and implementable strategy based on a few markers to grade TLS in three maturations stages of organization and functional significance. This transfer from high-level to few markers classification makes now feasible studies investigating the pathophysiological and medical relevance of grading, counting and locating TLS in pathological tissues, in digestive but also non-digestive organs as well as in cancer and non-cancer diseases.

## ACKNOWLEDGEMENTS

The authors would like to thank all the staff of the LBAI and the pathology department of CHU Brest for their help as well as the Brest Hospital Biobank BB-0033-00037. They also thank the Hyperion platform (LBAI, Brest, France) for its technical assistance.

## AUTHOR CONTRIBUTIONS

Involved in the conception or design of the work: MLR, JOP, AU. The conception of the panel: MLR, DB, PH, AU. Data acquisition: MLR, DB. Analysis of the data: MLR, PH, CLD. All contributing authors critically revised the article, contributed to the article, and approved the submitted version.

## SUPPLEMENTARY MATERIALS

**Supplementary Figure 1:**
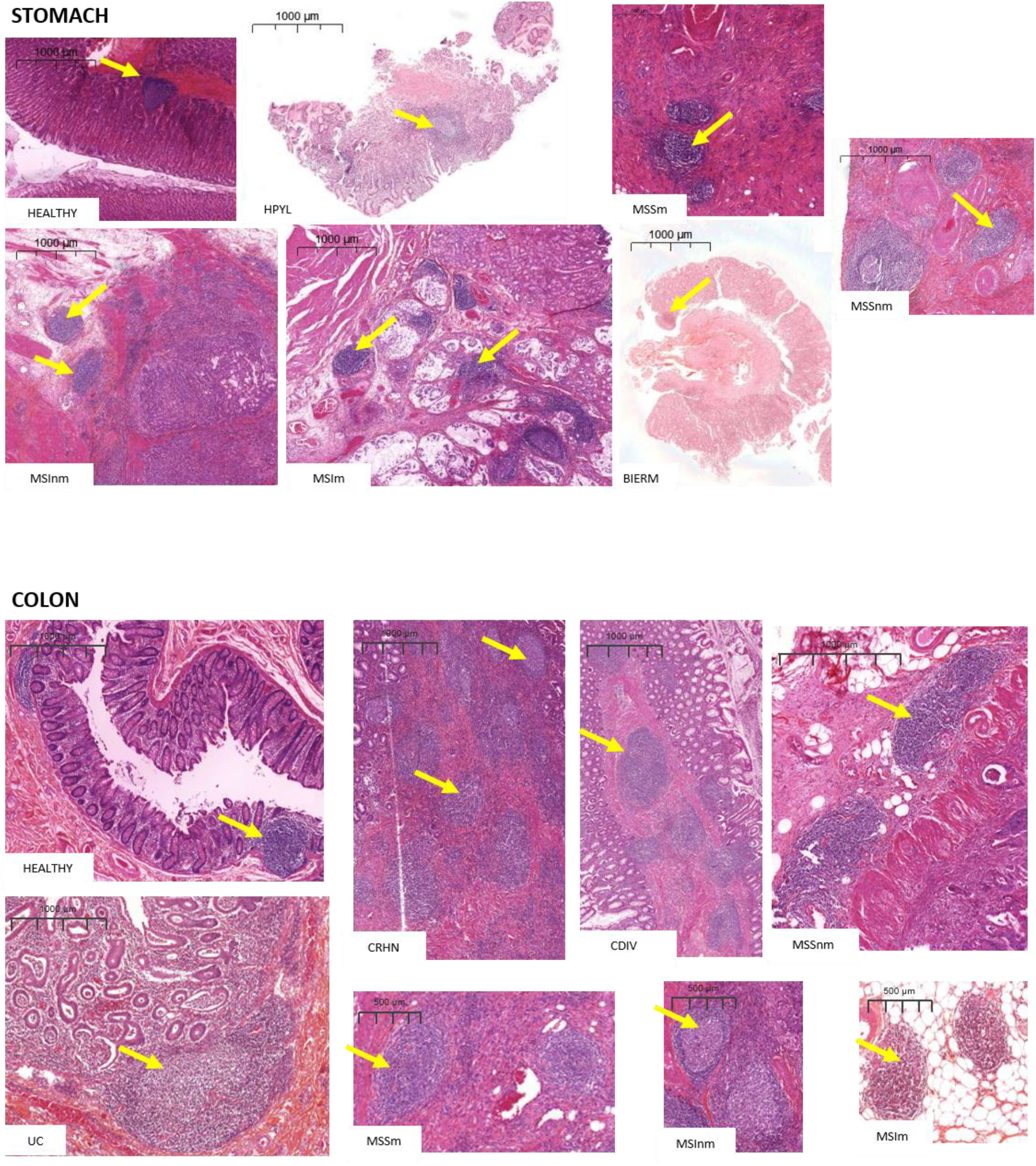
HES scans of the different conditions in the stomach and in the colon, yellow arrows indicate TLS observed and chosen by the pathologist.

**Supplementary Figure 2:**
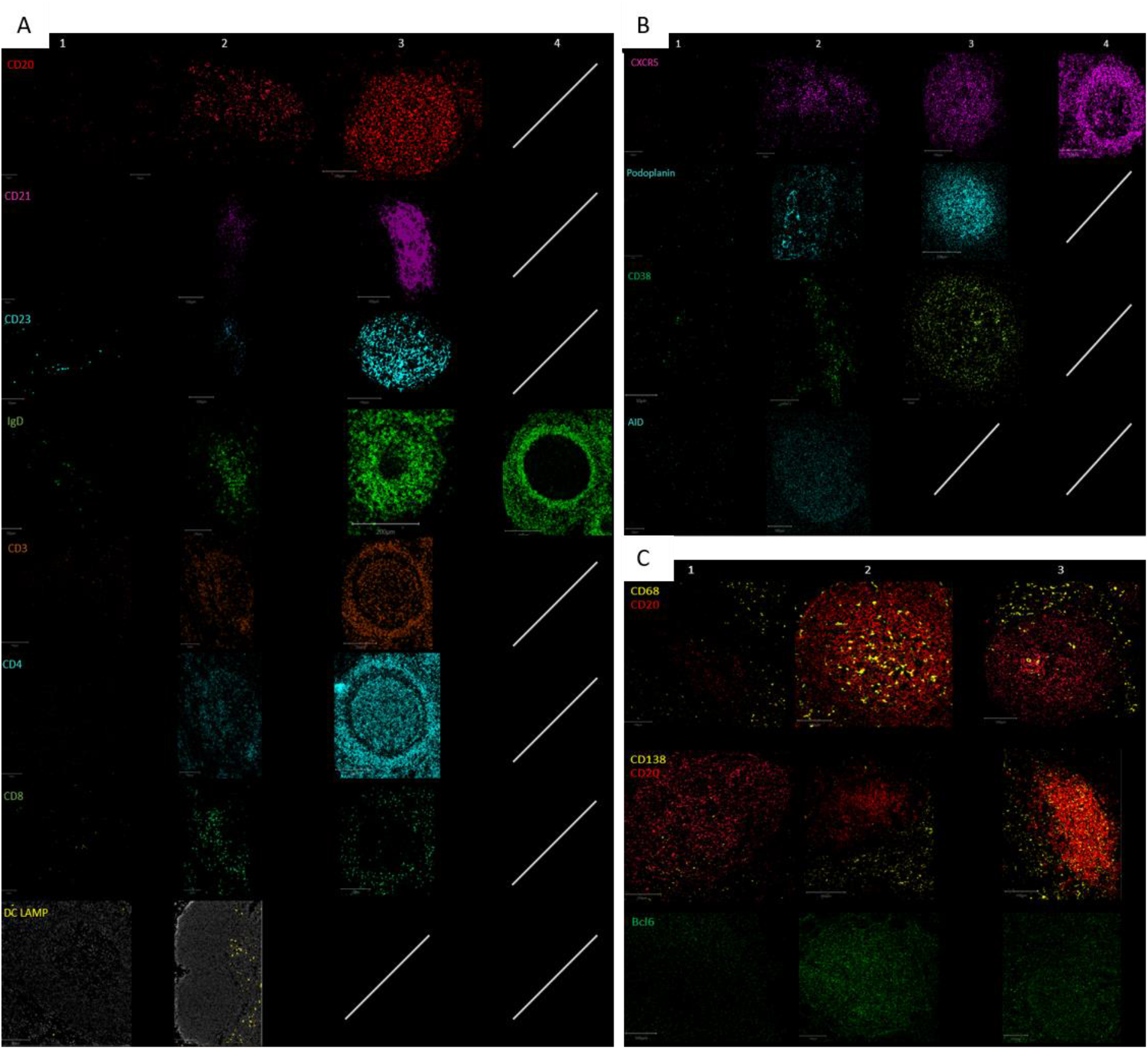
Grading of the morphological assessment of the TLS according to the morphological organization of (**A**) the 8 structural markers of a germinal center (**B**) the 4 functional markers (**C**) the 3 markers for the nodular interactions. The white oblique lines indicate that, this score does not exist for the marker considered.

**Supplementary Figure 3:**
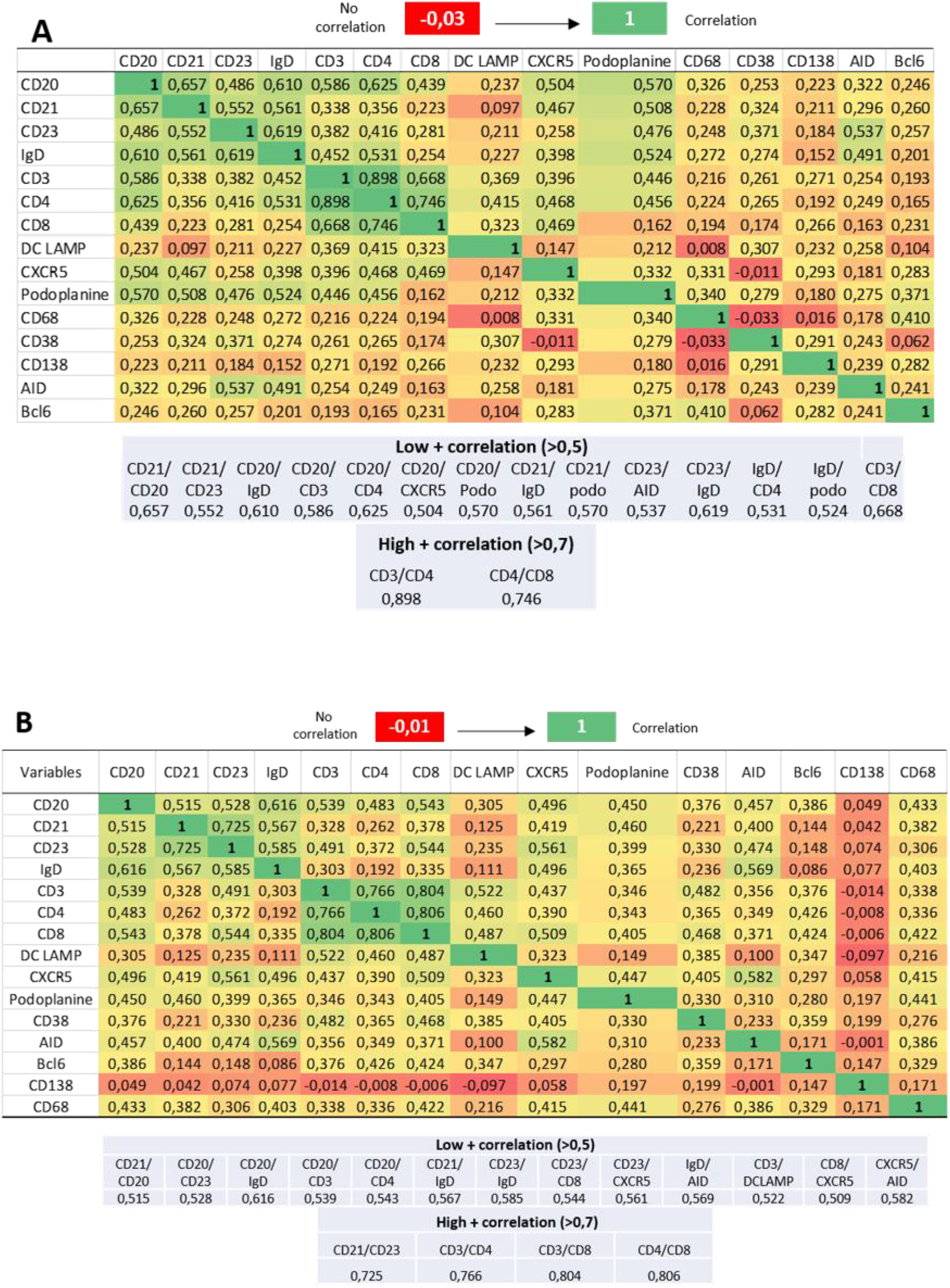
Matrix of the correlation scores of the 15 markers characterizing GC. Positive and negative correlations are shown in green and red, respectively **A**) In the stomach **B**) In the colon.

## BIBLIOGRAPHY

1. Tang H, Zhu M, Qiao J, et al. Lymphotoxin signalling in tertiary lymphoid structures and immunotherapy. Cell Mol Immunol 2017 Oct;14(10):809–18.

2. Aloisi F, Pujol-Borrell R. Lymphoid neogenesis in chronic inflammatory diseases. Nat Rev Immunol 2006 Mar;6(3):205–17.

3. Ansel KM, Cyster JG. Chemokines in lymphopoiesis and lymphoid organ development. Curr Opin Immunol 2001 Apr;13(2):172–9.

4. Dieu-Nosjean MC, Goc J, Giraldo NA, et al. Tertiary lymphoid structures in cancer and beyond. Trends Immunol 2014 Nov 1;35(11):571–80.

5. Schumacher TN, Thommen DS. Tertiary lymphoid structures in cancer. Science 2022 Jan 7;375(6576):eabf9419.

6. Calderaro J, Petitprez F, Becht E, et al. Intra-tumoral tertiary lymphoid structures are associated with a low risk of early recurrence of hepatocellular carcinoma. J Hepatol 2019 Jan;70(1):58–65.

7. Posch F, Silina K, Leibl S, et al. Maturation of tertiary lymphoid structures and recurrence of stage II and III colorectal cancer. Oncoimmunology 2018;7(2):e1378844.

8. Siliņa K, Soltermann A, Attar FM, et al. Germinal Centers Determine the Prognostic Relevance of Tertiary Lymphoid Structures and Are Impaired by Corticosteroids in Lung Squamous Cell Carcinoma. Cancer Res 2018 Mar 1;78(5):1308–20.

9. Winter S, Loddenkemper C, Aebischer A, et al. The chemokine receptor CXCR5 is pivotal for ectopic mucosa-associated lymphoid tissue neogenesis in chronic Helicobacter pylori-induced inflammation. J Mol Med 2010 Nov;88(11):1169–80.

10. Rugge M, Savarino E, Sbaraglia M, et al. Gastritis: The clinico-pathological spectrum. Dig Liver Dis Off J Ital Soc Gastroenterol Ital Assoc Study Liver 2021 Oct;53(10):1237–46.

11. Fabian O, Bajer L. Histopathological assessment of the microscopic activity in inflammatory bowel diseases: What are we looking for? World J Gastroenterol 2022 Sep 28;28(36):5300–12.

12. Pipi E, Nayar S, Gardner DH, et al. Tertiary Lymphoid Structures: Autoimmunity Goes Local. Front Immunol 2018;9:1952.

13. Sautès-Fridman C, Petitprez F, Calderaro J, et al. Tertiary lymphoid structures in the era of cancer immunotherapy. Nat Rev Cancer 2019 Jun;19(6):307–25.

14. Colbeck EJ, Ager A, Gallimore A, et al. Tertiary Lymphoid Structures in Cancer: Drivers of Antitumor Immunity, Immunosuppression, or Bystander Sentinels in Disease? Front Immunol 2017;8:1830.

15. Maoz A, Dennis M, Greenson JK. The Crohn’s-Like Lymphoid Reaction to Colorectal Cancer-Tertiary Lymphoid Structures With Immunologic and Potentially Therapeutic Relevance in Colorectal Cancer. Front Immunol 2019;10:1884.

16. Shokal U, Sharma PC. Implication of microsatellite instability in human gastric cancers. Indian J Med Res 2012 May;135(5):599–613.

17. Chang Q, Ornatsky OI, Siddiqui I, et al. Imaging Mass Cytometry. Cytometry A 2017;91(2):160–9.

18. Le Rochais M, Hemon P, Pers JO, Uguen A. Application of High-Throughput Imaging Mass Cytometry Hyperion in Cancer Research. Front Immunol 2022;13:859414.

19. Bankhead P, Loughrey MB, Fernández JA, et al. QuPath: Open source software for digital pathology image analysis. Sci Rep 2017 Dec;7(1):16878.

20. Dieu-Nosjean MC, Antoine M, Danel C, et al. Long-term survival for patients with non-small-cell lung cancer with intratumoral lymphoid structures. J Clin Oncol Off J Am Soc Clin Oncol 2008 Sep 20;26(27):4410–7.

21. Hiraoka N, Ino Y, Yamazaki-Itoh R, et al. Intratumoral tertiary lymphoid organ is a favourable prognosticator in patients with pancreatic cancer. Br J Cancer 2015 May 26;112(11):1782–90.

22. Siliņa K, Rulle U, Kalniņa Z, et al. Manipulation of tumour-infiltrating B cells and tertiary lymphoid structures: a novel anti-cancer treatment avenue? Cancer Immunol Immunother Cell 2014 Jul;63(7):643–62.

23. Fridman WH, Zitvogel L, Sautès-Fridman C, et al. The immune contexture in cancer prognosis and treatment. Nat Rev Clin Oncol 2017 Dec;14(12):717–34.

24. Helmink BA, Reddy SM, Gao J, et al. B cells and tertiary lymphoid structures promote immunotherapy response. Nature 2020 Jan;577(7791):549–55.

25. Cabrita R, Lauss M, Sanna A, et al. Tertiary lymphoid structures improve immunotherapy and survival in melanoma. Nature 2020 Jan;577(7791):561–5.

26. Petitprez F, de Reyniès A, Keung EZ, et al. B cells are associated with survival and immunotherapy response in sarcoma. Nature 2020 Jan;577(7791):556–60.

