## Supplementary Figure 1 for "Deciphering the maturation of tertiary lymphoid structures in cancer and inflammatory diseases of the digestive tract using imaging mass cytometry: from high-level data to a simple architectural and functional grading"

**Supplementary Figure 1:** HES scans of the different conditions in the stomach and in the colon, yellow arrows indicate TLS observed and chosen by the pathologist.

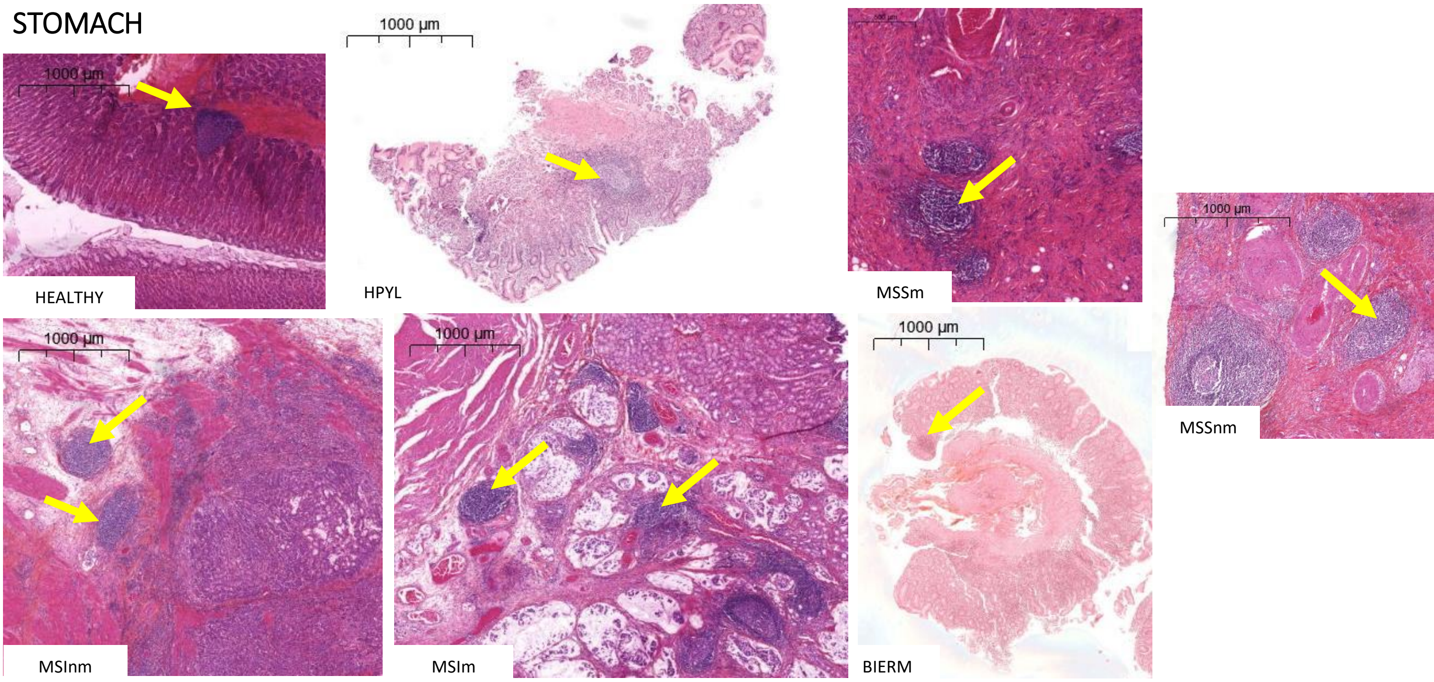


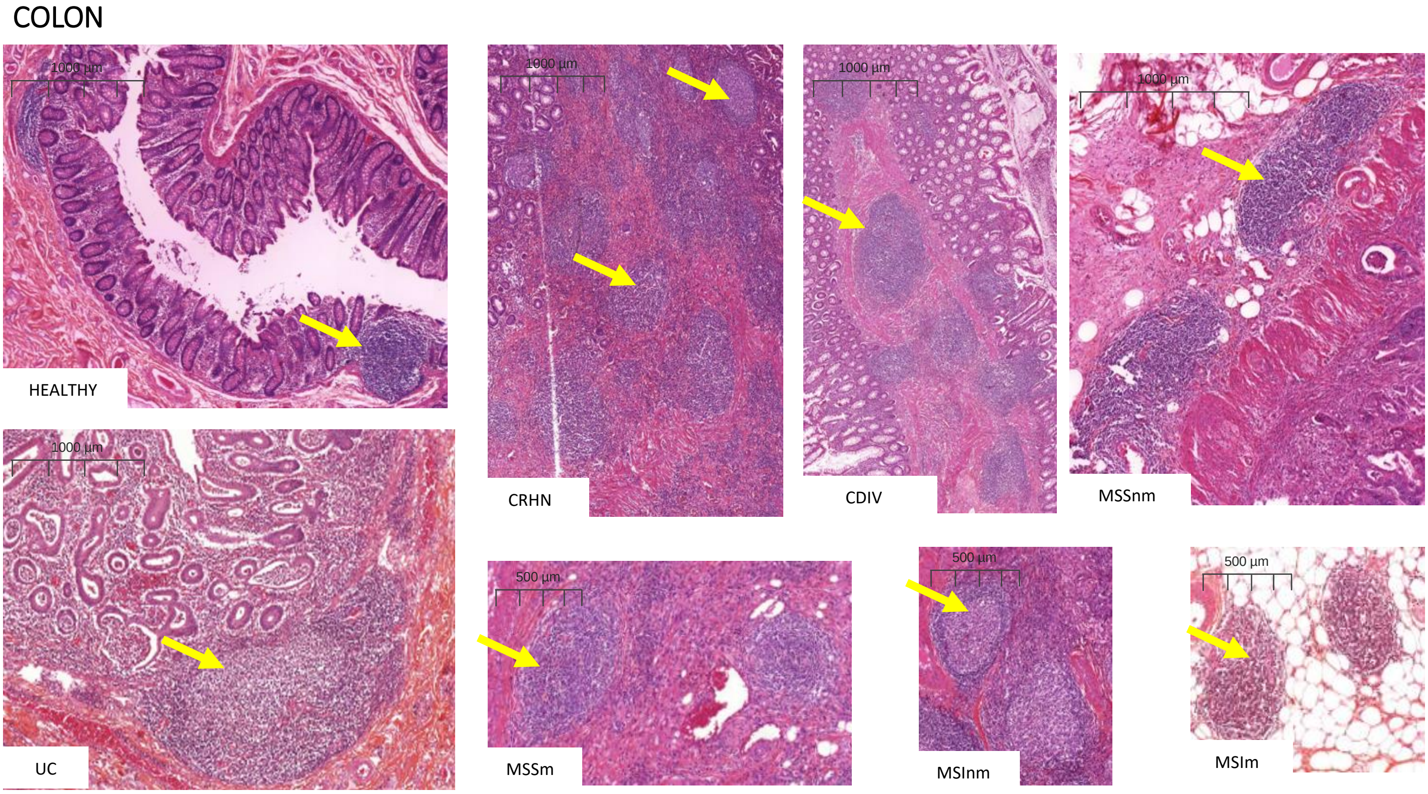
