## Supplementary Figure 2 for "Deciphering the maturation of tertiary lymphoid structures in cancer and inflammatory diseases of the digestive tract using imaging mass cytometry: from high-level data to a simple architectural and functional grading"

**Supplementary Figure 2:** Grading of the morphological assessment of the TLS according to the morphological organization of (**A**) the 8 structural markers of a germinal center (**B**) the 4 functional markers (**C**) the 3 markers for the nodular interactions. The white oblique lines indicate that, this score do not exist for the marker considered.

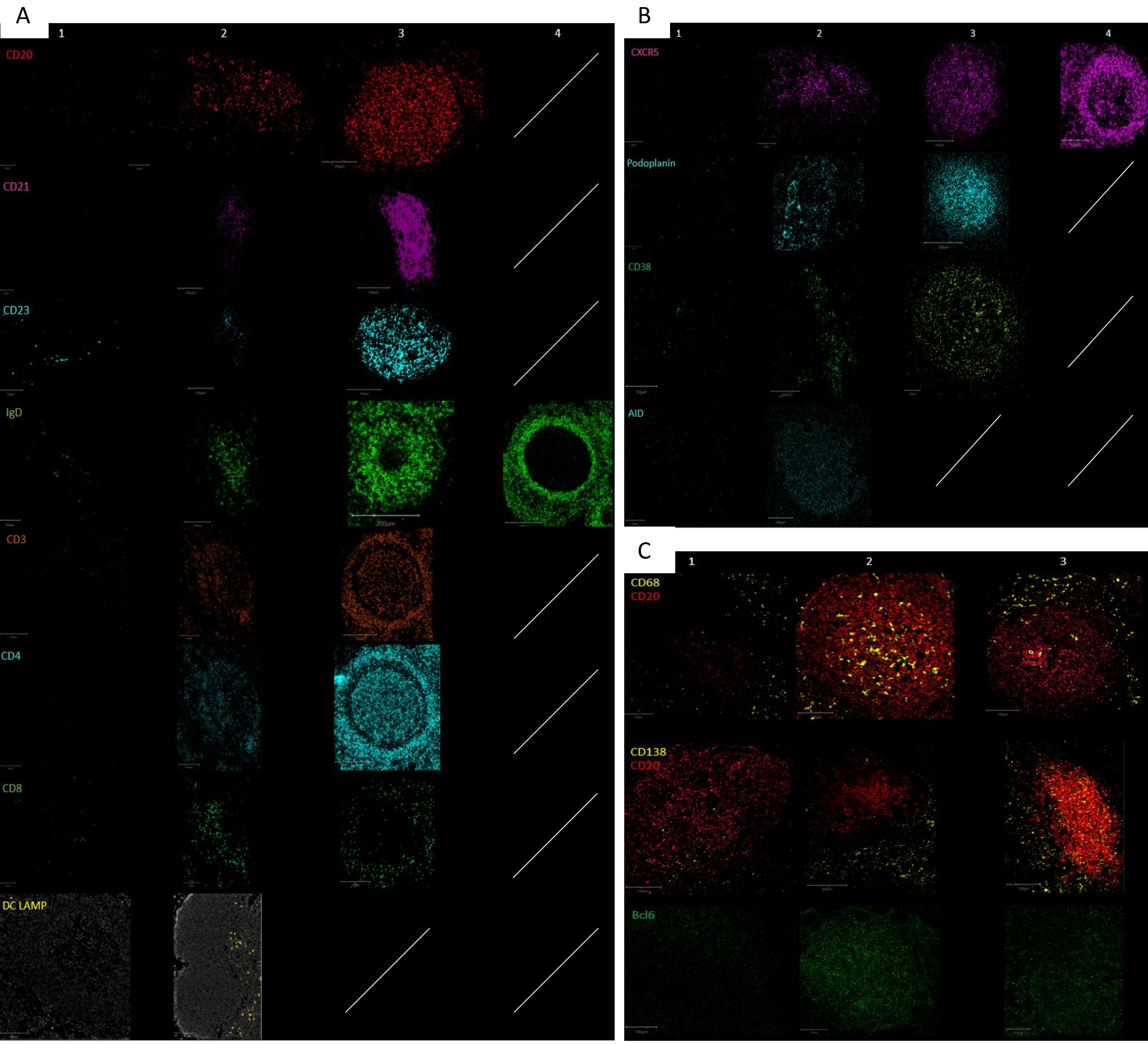
