## Supplementary Figure 3 for "Deciphering the maturation of tertiary lymphoid structures in cancer and inflammatory diseases of the digestive tract using imaging mass cytometry: from high-level data to a simple architectural and functional grading"

**Supplementary Figure 3:** Matrix of the correlation scores of the 15 markers characterizing GC. Positive and negative correlations are shown in green and red, respectively **A**) In the stomach **B**) In the colon.

**A**

No correlation -0,03 → 1 Correlation

| | CD20 | CD21 | CD23 | IgD | CD3 | CD4 | CD8 | DC LAMP | CXCR5 | Podoplanine | CD68 | CD38 | CD138 | AID | Bcl6 |
|---|---|---|---|---|---|---|---|---|---|---|---|---|---|---|---|
| CD20 | 1 | 0,657 | 0,486 | 0,610 | 0,586 | 0,625 | 0,439 | 0,237 | 0,504 | 0,570 | 0,326 | 0,253 | 0,223 | 0,322 | 0,246 |
| CD21 | 0,657 | 1 | 0,552 | 0,561 | 0,338 | 0,356 | 0,223 | 0,097 | 0,467 | 0,508 | 0,228 | 0,324 | 0,211 | 0,296 | 0,260 |
| CD23 | 0,486 | 0,552 | 1 | 0,619 | 0,382 | 0,416 | 0,281 | 0,211 | 0,258 | 0,476 | 0,248 | 0,371 | 0,184 | 0,537 | 0,257 |
| IgD | 0,610 | 0,561 | 0,619 | 1 | 0,452 | 0,531 | 0,254 | 0,227 | 0,398 | 0,524 | 0,272 | 0,274 | 0,152 | 0,491 | 0,201 |
| CD3 | 0,586 | 0,338 | 0,382 | 0,452 | 1 | 0,898 | 0,668 | 0,369 | 0,396 | 0,446 | 0,216 | 0,261 | 0,271 | 0,254 | 0,193 |
| CD4 | 0,625 | 0,356 | 0,416 | 0,531 | 0,898 | 1 | 0,746 | 0,415 | 0,468 | 0,456 | 0,224 | 0,265 | 0,192 | 0,249 | 0,165 |
| CD8 | 0,439 | 0,223 | 0,281 | 0,254 | 0,668 | 0,746 | 1 | 0,323 | 0,469 | 0,162 | 0,194 | 0,174 | 0,266 | 0,163 | 0,231 |
| DC LAMP | 0,237 | 0,097 | 0,211 | 0,227 | 0,369 | 0,415 | 0,323 | 1 | 0,147 | 0,212 | 0,008 | 0,307 | 0,232 | 0,258 | 0,104 |
| CXCR5 | 0,504 | 0,467 | 0,258 | 0,398 | 0,396 | 0,468 | 0,469 | 0,147 | 1 | 0,332 | 0,331 | -0,011 | 0,293 | 0,181 | 0,283 |
| Podoplanine | 0,570 | 0,508 | 0,476 | 0,524 | 0,446 | 0,456 | 0,162 | 0,212 | 0,332 | 1 | 0,340 | 0,279 | 0,180 | 0,275 | 0,371 |
| CD68 | 0,326 | 0,228 | 0,248 | 0,272 | 0,216 | 0,224 | 0,194 | 0,008 | 0,331 | 0,340 | 1 | -0,033 | 0,016 | 0,178 | 0,410 |
| CD38 | 0,253 | 0,324 | 0,371 | 0,274 | 0,261 | 0,265 | 0,174 | 0,307 | -0,011 | 0,279 | -0,033 | 1 | 0,291 | 0,243 | 0,062 |
| CD138 | 0,223 | 0,211 | 0,184 | 0,152 | 0,271 | 0,192 | 0,266 | 0,232 | 0,293 | 0,180 | 0,016 | 0,291 | 1 | 0,239 | 0,282 |
| AID | 0,322 | 0,296 | 0,537 | 0,491 | 0,254 | 0,249 | 0,163 | 0,258 | 0,181 | 0,275 | 0,178 | 0,243 | 0,239 | 1 | 0,241 |
| Bcl6 | 0,246 | 0,260 | 0,257 | 0,201 | 0,193 | 0,165 | 0,231 | 0,104 | 0,283 | 0,371 | 0,410 | 0,062 | 0,282 | 0,241 | 1 |

**Low + correlation (>0,5)**

| CD21/ CD20 | CD21/ CD23 | CD20/ IgD | CD20/ CD3 | CD20/ CD4 | CD20/ CXCR5 | CD20/ Podo | CD21/ IgD | CD21/ podo | CD23/ AID | CD23/ IgD | IgD/ CD4 | IgD/ podo | CD3/ CD8 |
|---|---|---|---|---|---|---|---|---|---|---|---|---|---|
| 0,657 | 0,552 | 0,610 | 0,586 | 0,625 | 0,504 | 0,570 | 0,561 | 0,570 | 0,537 | 0,619 | 0,531 | 0,524 | 0,668 |

**High + correlation (>0,7)**

| CD3/CD4 | CD4/CD8 |
|---|---|
| 0,898 | 0,746 |

**B**

No correlation -0,01 → 1 Correlation

| Variables | CD20 | CD21 | CD23 | IgD | CD3 | CD4 | CD8 | DC LAMP | CXCR5 | Podoplanine | CD38 | AID | Bcl6 | CD138 | CD68 |
|---|---|---|---|---|---|---|---|---|---|---|---|---|---|---|---|
| CD20 | 1 | 0,515 | 0,528 | 0,616 | 0,539 | 0,483 | 0,543 | 0,305 | 0,496 | 0,450 | 0,376 | 0,457 | 0,386 | 0,049 | 0,433 |
| CD21 | 0,515 | 1 | 0,725 | 0,567 | 0,328 | 0,262 | 0,378 | 0,125 | 0,419 | 0,460 | 0,221 | 0,400 | 0,144 | 0,042 | 0,382 |
| CD23 | 0,528 | 0,725 | 1 | 0,585 | 0,491 | 0,372 | 0,544 | 0,235 | 0,561 | 0,399 | 0,330 | 0,474 | 0,148 | 0,074 | 0,306 |
| IgD | 0,616 | 0,567 | 0,585 | 1 | 0,303 | 0,192 | 0,335 | 0,111 | 0,496 | 0,365 | 0,236 | 0,569 | 0,086 | 0,077 | 0,403 |
| CD3 | 0,539 | 0,328 | 0,491 | 0,303 | 1 | 0,766 | 0,804 | 0,522 | 0,437 | 0,346 | 0,482 | 0,356 | 0,376 | -0,014 | 0,338 |
| CD4 | 0,483 | 0,262 | 0,372 | 0,192 | 0,766 | 1 | 0,806 | 0,460 | 0,390 | 0,343 | 0,365 | 0,349 | 0,426 | -0,008 | 0,336 |
| CD8 | 0,543 | 0,378 | 0,544 | 0,335 | 0,804 | 0,806 | 1 | 0,487 | 0,509 | 0,405 | 0,468 | 0,371 | 0,424 | -0,006 | 0,422 |
| DC LAMP | 0,305 | 0,125 | 0,235 | 0,111 | 0,522 | 0,460 | 0,487 | 1 | 0,323 | 0,149 | 0,385 | 0,100 | 0,347 | -0,097 | 0,216 |
| CXCR5 | 0,496 | 0,419 | 0,561 | 0,496 | 0,437 | 0,390 | 0,509 | 0,323 | 1 | 0,447 | 0,405 | 0,582 | 0,297 | 0,058 | 0,415 |
| Podoplanine | 0,450 | 0,460 | 0,399 | 0,365 | 0,346 | 0,343 | 0,405 | 0,149 | 0,447 | 1 | 0,330 | 0,310 | 0,280 | 0,197 | 0,441 |
| CD38 | 0,376 | 0,221 | 0,330 | 0,236 | 0,482 | 0,365 | 0,468 | 0,385 | 0,405 | 0,330 | 1 | 0,233 | 0,359 | 0,199 | 0,276 |
| AID | 0,457 | 0,400 | 0,474 | 0,569 | 0,356 | 0,349 | 0,371 | 0,100 | 0,582 | 0,310 | 0,233 | 1 | 0,171 | -0,001 | 0,386 |
| Bcl6 | 0,386 | 0,144 | 0,148 | 0,086 | 0,376 | 0,426 | 0,424 | 0,347 | 0,297 | 0,280 | 0,359 | 0,171 | 1 | 0,147 | 0,329 |
| CD138 | 0,049 | 0,042 | 0,074 | 0,077 | -0,014 | -0,008 | -0,006 | -0,097 | 0,058 | 0,197 | 0,199 | -0,001 | 0,147 | 1 | 0,171 |
| CD68 | 0,433 | 0,382 | 0,306 | 0,403 | 0,338 | 0,336 | 0,422 | 0,216 | 0,415 | 0,441 | 0,276 | 0,386 | 0,329 | 0,171 | 1 |

**Low + correlation (>0,5)**

| CD21/ CD20 | CD20/ CD23 | CD20/ IgD | CD20/ CD3 | CD20/ CD4 | CD21/ IgD | CD23/ IgD | CD23/ CD8 | CD23/ CXCR5 | IgD/ AID | CD3/ DCLAMP | CD8/ CXCR5 | CXCR5/ AID |
|---|---|---|---|---|---|---|---|---|---|---|---|---|
| 0,515 | 0,528 | 0,616 | 0,539 | 0,543 | 0,567 | 0,585 | 0,544 | 0,561 | 0,569 | 0,522 | 0,509 | 0,582 |

**High + correlation (>0,7)**

| CD21/CD23 | CD3/CD4 | CD3/CD8 | CD4/CD8 |
|---|---|---|---|
| 0,725 | 0,766 | 0,804 | 0,806 |
